# The cytokinetic midbody mediates asymmetric fate specification at mitotic exit during neural stem cell division

**DOI:** 10.1101/2024.08.27.609974

**Authors:** Bryce LaFoya, Rhiannon R Penkert, Kenneth E. Prehoda

## Abstract

Asymmetric cell division (ACD) is a broadly used mechanism for generating cellular diversity. Molecules known as fate determinants are segregated during ACD to generate distinct sibling cell fates, but determinants should not be activated until fate can be specified asymmetrically. Determinants could be activated after cell division but many animal cells complete division long after mitosis ends, raising the question of how activation could occur at mitotic exit taking advantage of the unique state plasticity at this time point. Here we show that the midbody, a microtubule-rich structure that forms in the intercellular bridge connecting nascent siblings, mediates fate determinant activation at mitotic exit in neural stem cells (NSCs) of the *Drosophila* larval brain. The fate determinants Prospero (Pros) and Brain tumor (Brat) are sequestered at the NSC membrane at metaphase but are released immediately following nuclear division when the midbody forms, well before cell division completes. The midbody isolates nascent sibling cytoplasms, allowing determinant release from the membrane via the cell cycle phosphatase String, without influencing the fate of the incorrect sibling. Our results identify the midbody as a key facilitator of ACD that allows asymmetric fate determinant activation to be initiated before division.

## Introduction

Asymmetric cell division (ACD) is a fundamental mechanism for generating cellular diversity, employed by organisms across the tree of life^1–6^. During development and homeostasis, ACDs generate specialized cell types by producing sibling cells that assume distinct fates. Factors known as fate determinants are central players in ACDs as they maintain or alter sibling cell state. The segregation of fate determinants into the siblings, through determinant polarization and division plane alignment, is the central feature of current ACD models^2^. However, segregation may not fully capture the essential features of ACD. Transcription factor fate determinant activation is an example of an essential ACD process that is not explained by segregation. Many fate determinants are transcription factors that are polarized on the membrane during division to facilitate segregation but must be released from the membrane and imported into the nucleus to regulate gene activity. This example highlights a critical knowledge gap in our understanding of ACD: how fate determinants are activated to achieve asymmetric sibling cell states.

The model for intrinsic ACDs (those that are cell autonomous) has been well-established for at least 30 years^1^ but doesn’t account for fate determinant regulation. In the ACD standard model, segregation of fate determinants is the central feature responsible for the distinct fates of the resulting sibling cells. Fate determinants are segregated by their polarization during the division process and alignment of the polarity and division axes, causing their specific deposition into only one sibling^1,2,4,5,7^. While segregation determines which sibling receives a determinant, it does not specify whether the determinant is active in the sibling. In fact, for reasons we outline here, fate determinant activity transitions are likely general features of ACD that are hidden within the current segregation-focused standard model (Fig. 1A). These transitions are likely to be highly regulated since the activity of determinants before sibling separation could cause the asymmetry of the division to be lost. The fate determinant Prospero (Pros in *Drosophila*; Prox1 in mammals) functions in asymmetrically dividing *Drosophila* neural stem cells (NSCs; aka neuroblasts) and illustrates this problem^8–11^. Pros is a transcription factor that is critical for specifying the fate of the differentiating sibling cell (NP; neural progenitor) following an NSC division (the other sibling remains an NSC; Fig. 1A). During division, Pros is polarized to the basal plasma membrane but eventually becomes nuclear in the NP sibling where it can bind chromatin and alter gene expression^8,9^. Thus, Pros’ membrane binding plays a dual role – along with the commonly understood function to facilitate segregation, it also inhibits Pros’ transcription factor activity by sequestering it away from the nucleus. Pros must ultimately be released from the membrane, but only when doing so would allow the NP fate to be specified without corrupting the fate of the NSC sibling. Understanding when and how Pros is activated is therefore a critical aspect of the NSC ACD mechanism, but little is known about this essential transition. In terms of the standard ACD model, the Pros example highlights what has been an implicit but essential feature of the model – that fate determinants are inactive during the polarization step but become activated at some later point.

**Figure 1.**
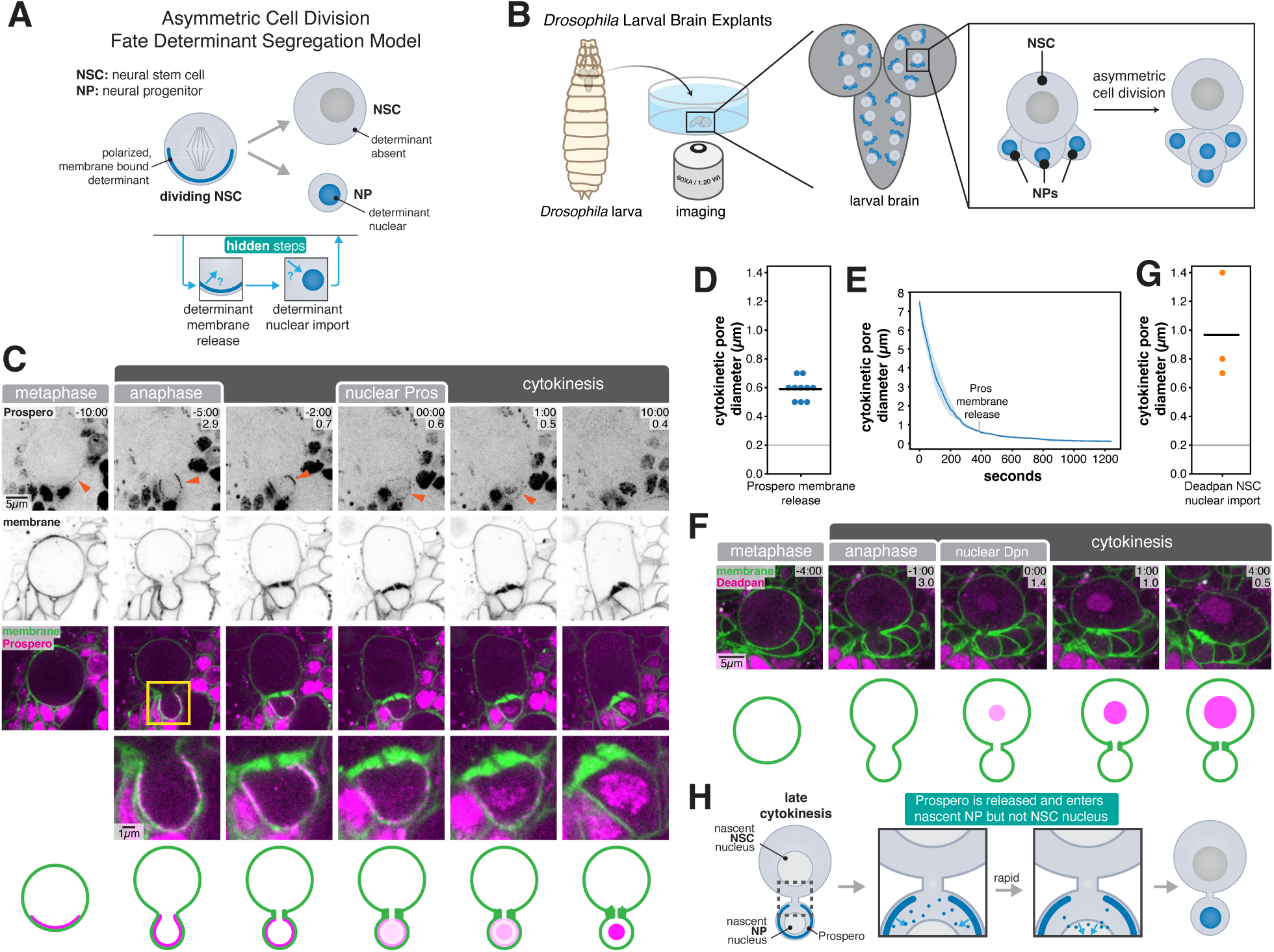
Asymmetric fate specification is initiated before NSC division completes. (A) Uncovering the hidden steps in asymmetric cell division. The standard model (top) focuses on fate determinant segregation through polarization and division plane orientation. The focus on segregation obscures a key element of the model that is likely a general feature, fate determinant regulation (bottom). In this example, the fate determinant is a transcription factor polarized on the plasma membrane during neural stem cell (NSC) division. The transcription factor must be released from the membrane and imported into the nucleus to influence the neural progenitor (NP) sibling state. (B) Imaging NSC asymmetric divisions. Larval Drosophila brains explants are cultured and imaged using spinning disk confocal microscopy. NSCs and their progeny are identified through the position in the brain and expression of membrane marker driven by the NSC driver, Worniu-GAL4. (C) Dynamics of the fate determinant Prospero (Pros) during the late stages of NSC asymmetric division. Frames from Video 1 are shown with Pros-GFP expressed from its endogenous locus and the plasma membrane marker PLCδ-PH-mCherry (expressed with Worniu-GAL4 driven UAS; “membrane”) through an optical section containing the cytokinetic pore. Arrowheads point to cortical Pros signal in dividing NSC and nascent NP sibling. Time in minutes relative to the presence of nuclear Pros is shown along with the diameter of the cytokinetic pore in microns. Panels containing both Pros and membrane signal are shown in the third row along with an inset in the fourth row that focuses on the cytokinetic pore and NP sibling. Bottom row shows a schematized representation of the localization within the dividing NSC. (D) Quantification of Pros membrane release. The diameters of the pore connecting nascent siblings when nuclear signal was detected for Pros is shown. Each point represents a distinct NSC division. Abscission, the final step of cytokinesis, does not occur until sometime after the pore reaches 200 nm (grey line). (E) Quantification of cytokinetic pore closure dynamics during NSC ACD. The solid line represents the mean of five distinct NSC divisions and the shaded region represents one standard deviation. (F) Dynamics of asymmetric nuclear Deadpan (Dpn). Frames from Video 1 are shown with Dpn-GFP expressed from its endogenous locus and the plasma membrane marker PLCδ-PH-mCherry (expressed with Worniu-GAL4 driven UAS) through a section containing the cytokinetic pore. Time in minutes relative to presence of nuclear Dpn is shown along with the diameter of the cytokinetic pore in microns. (G) Quantification of Dpn nuclear import. The diameters of the pore connecting nascent siblings when nuclear signal was detected for Dpn are shown as in (D). (H) Pre-division initiation of asymmetric fate specification. Pros is released from the membrane and enters the nascent NP nucleus without entering the NSC nucleus.

Cell fate transitions have been proposed to be coupled to cell division^6,12,13^ suggesting that fate determinant activation could be linked to the final step of division, abscission. Division (i.e. cytokinesis) is a seemingly natural boundary for determinant activation and asymmetric fate specification because determinants cannot pass between siblings following abscission. On the other hand, transcriptional programs that control cellular identity can be rapidly set at mitotic exit because of chromosome decompaction and epigentic changes^14–17^, including global reactivation of transcription^18^. However, many animal cells undergo abscission long after mitosis with the nascent sibling cells remaining connected by an intercellular bridge for an extended period in G1 phase^19–21^. Thus, the delay between the end of mitosis and cell division represents a paradox – initiating fate determinant activation after abscission would safely allow for asymmetric fate specification but with the ideal window for altering cell fate potentially having passed. We sought to resolve this paradox and determine when fate determinant activation is initiated by examining determinant and cell cycle dynamics in the asymmetrically dividing NSCs of the *Drosophila* larval brain.

## Results

### Fate determinant activation begins before cell division completes

The fate determinant Pros localizes to the NSC basal membrane at metaphase^8,9^. Pros is ultimately released from the membrane so that it can enter the nucleus of the NP sibling. A previous study examined Pros nuclear import relative to early furrowing^22^ but the timing of release and nuclear entry relative to the late steps of cytokinesis has not been known. We used high speed, super-resolution live imaging NSCs (Fig. 1B) expressing Pros-GFP and the membrane marker PLCδ-PH-mCherry, to determine when Pros membrane release occurs relative to cytokinesis. Pros was initially targeted to the basal membrane several minutes before furrow ingression began where it remained until late cytokinesis (Fig. 1C,D and Video 1). Surprisingly, Pros was released from the membrane while the nascent siblings remained connected by the intercellular bridge, a thin, pore-forming tube of plasma membrane. We measured an average cytokinetic pore diameter of approximately 0.6 µm when Pros was released from the membrane (0.59 ± 0.07 µm; 1 SD, n = 10 divisions from distinct NSCs). Pros rapidly entered the nucleus after release without any significant cytoplasmic accumulation. Pore constriction in late cytokinesis is slow in the dividing NSC^23^ (Fig. 1E), leading to a substantial delay between Pros membrane release and the pore reaching the diameter at which abscission can take place (approximately 200 nm)^24^. We also observed large tubules or sheets of plasma membrane forming near the pore during this phase (Fig. 1C and Video 1), consistent with a previous report^25^. We conclude that the initial Pros activating step – its release from the membrane – and its subsequent entry into the NP nucleus, occur well before cell division completes.

The timing of Pros translocation from the membrane to the nascent NP nucleus before division prompted us to examine fate determinant activation in the nascent NSC. Deadpan (Dpn) is a transcription factor that specifies NSC fate^26^ and localizes specifically to the NSC nucleus. We found that Dpn entered the nascent NSC nucleus before division completed (Fig. 1F,G; pore diameter 0.97 ± 0.38 µm; 1 SD, n = 3 divisions from distinct NSCs), on average slightly earlier than Pros entered the nascent NP nucleus. Thus, asymmetric fate specification is initiated in both the nascent NSC and NP siblings before cell division completes (Fig. 1H).

Asymmetric fate specification likely occurs after nuclear division (i.e. mitosis) completes, and the rapid entry of Pros into the nascent NP nucleus immediately following membrane release is consistent with this hypothesis. We imaged several markers for the end of mitosis along with the plasma membrane to compare the relative timing of nuclear and cell division. Mitotic chromosomes were decompacted at a pore size of approximately 3 µm (3.0 ± 0.7; n = 3), as assessed by Histone 2A (Fig. 2A,B; Video 2). Nuclei were separated at 1.3 ± 0.1 µm (n = 3) pore size according to the nuclear membrane marker Klaroid (Fig. 2B,C; Video 2). Finally, an NLS-DsRed fusion was imported into the nascent NSC and NP nuclei at 1.0 ± 0.4 µm (n = 3) pore size (Fig. 2B,D; Video 2). Together, these observations establish a timeline of events that occur during late mitosis and cytokinesis, along with the initiation of asymmetric fate determinant activation (Fig. 2E). Fate determinant activation is initiated immediately following the events of late mitosis, but well before the cytokinetic pore reaches 200 nm and the ESCRT-III machinery can carry out the final step of cytokinesis, the resolution of the plasma membrane or abscission^24^. The entry of fate determinants into the nucleus before abscission suggests that a mechanism exists that allows asymmetric fate specification to occur while the nascent siblings remain connected.

**Figure 2.**
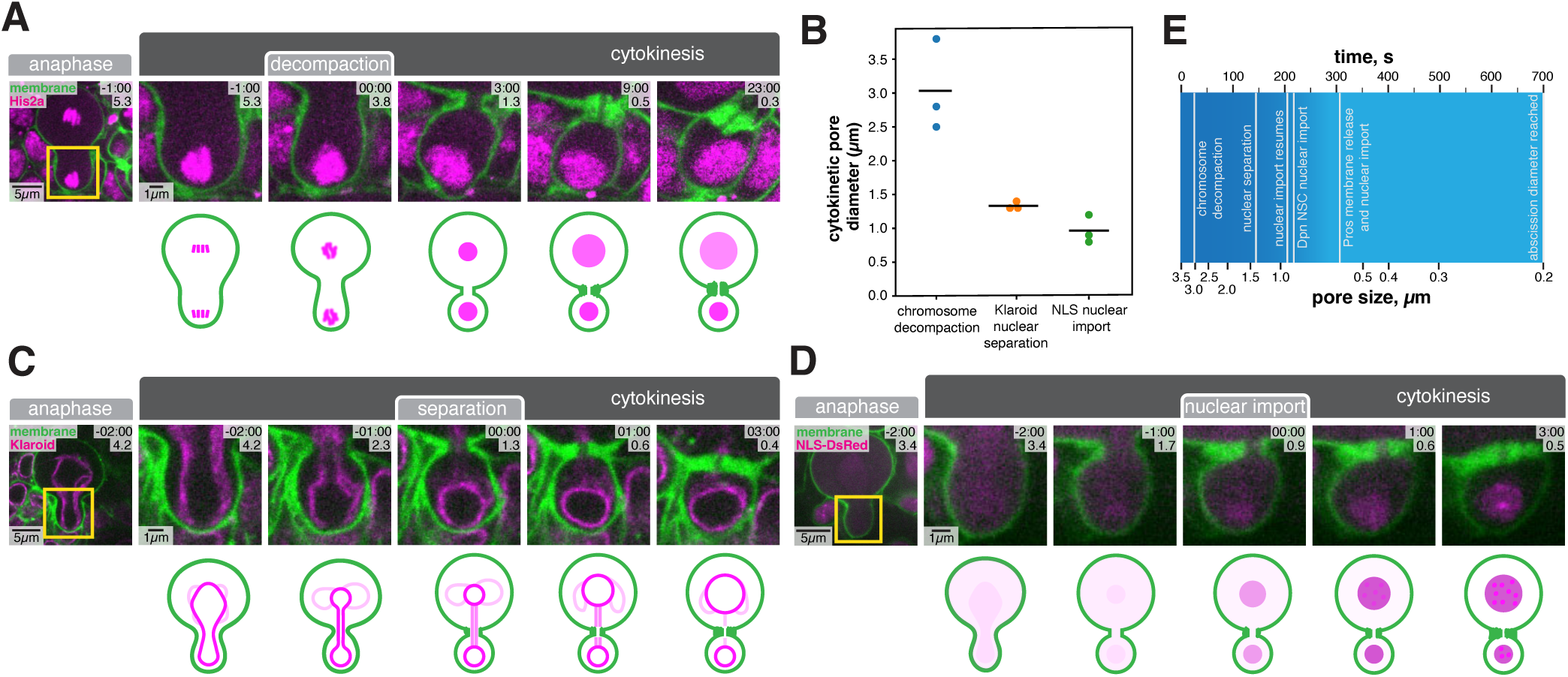
A temporal window between mitosis and cytokinesis during NSC ACD. (A) Chromatin dynamics during late NSC division focusing on the cytokinetic pore and nascent NP sibling. Frames from Video 2 are shown with PLCδ-PH-GFP (expressed with Worniu-GAL4 driven UAS) and the chromatin marker RFP tagged Histone 2A (His2a-RFP expressed from its endogenous locus) through an optical section containing the cytokinetic pore. Time in minutes relative to the chromosome decompaction is shown along with the diameter of the cytokinetic pore in microns. A schematic representation of the localization within the dividing NSC is shown below. (B) Quantification of chromosome decompaction, nuclear separation, and nuclear import, relative to cytokinetic pore diameter. Each point represents a distinct NSC division. (C) Nuclear dynamics during late NSC division focusing on the cytokinetic pore and nascent NP sibling. Frames from Video 2 are shown with Klaroid-GFP expressed from its native promoter and the plasma membrane marker PLCδ-PH-mCherry (expressed with worniu-GAL4 driven UAS) through an optical section containing the cytokinetic pore. Arrowheads point to cortical Pros signal in dividing NSC and nascent NP sibling. Time in minutes relative to nuclear membrane separation is shown along with the diameter of the cytokinetic pore in microns. A schematic representation of the nascent siblings is shown below. (D) Nuclear import dynamics during late NSC division focusing on the cytokinetic pore and nascent NP sibling. Frames from Video 2 are shown with PLCδ-PH-GFP and NLS-DsRed (both expressed with Worniu-GAL4 driven UAS) through an optical section containing the cytokinetic pore. Time in minutes relative to nuclear import onset is shown along with the diameter of the cytokinetic pore in microns. A schematic representation of the nascent siblings is shown below. (E) Timeline of late NSC division and initiation of fate determinant activation.

### Nascent sibling cytoplasms are isolated before cell division completes

Fate determinant release into the cytoplasm while the nascent siblings remain connected raises the question of how determinants are prevented from corrupting the fate of the incorrect sibling (e.g. why doesn’t Pros enter the nascent NSC cytoplasm and nucleus?). One possibility is that cellular structures besides the plasma membrane prevent fate determinants from escaping their nascent sibling’s cytoplasm after membrane release, similar to the function of the bud neck in budding yeast^27,28^. To test this hypothesis, we examined the dynamics of the fate determinant Brain tumor (Brat). Brat is a translational repressor that functions in the cytoplasm of the NP once it is released from the membrane^29,30^. We used Brat’s cytoplasmic localization to determine whether cytoplasmic exchange occurs between the nascent siblings. Brat was released from the membrane when the cytokinetic pore reached a diameter of approximately 0.7 µm (Fig. 3A,B; Video 3), similar to when Pros is released. There was a correspondingly rapid increase in cytoplasmic Brat that was specific to the nascent NP (Fig. 3B). Brat’s accumulation in the nascent NP cytoplasm indicates that the pore connecting the nascent siblings impedes Brat transfer into the nascent NSC cytoplasm.

**Figure 3.**
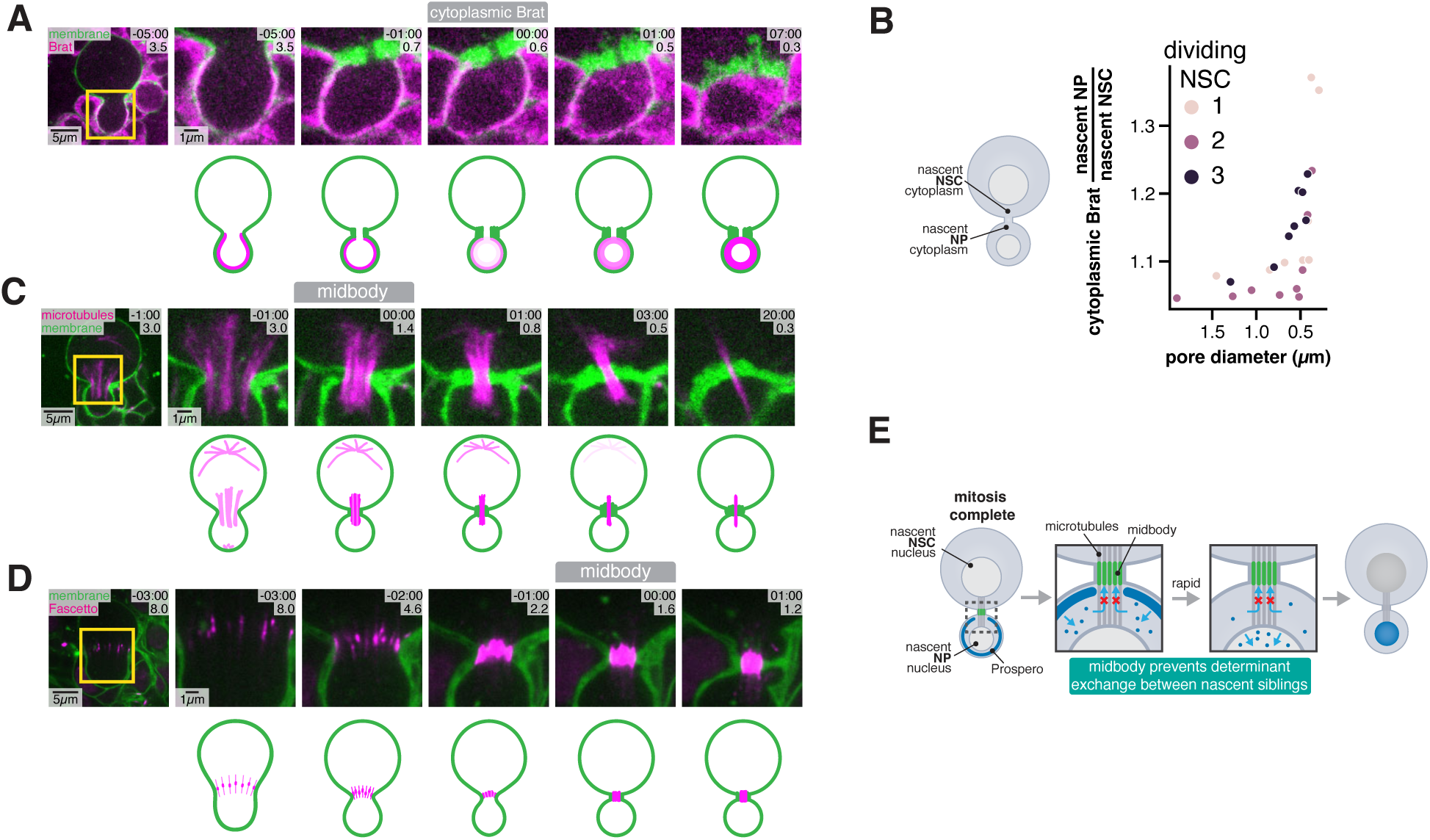
Fate determinants are isolated in nascent siblings before abscission. (A) Dynamics of the fate determinant Brain tumor (Brat) during the late stages of NSC asymmetric division focusing on the cytokinetic pore and nascent NP sibling. Frames from Video 3 are shown with GFP-Brat expressed from its endogenous locus and the plasma membrane marker PLCδ-PH-mCherry (expressed with Worniu-GAL4 driven UAS) through an optical section containing the cytokinetic pore. Time in minutes relative to Brat translocation from the membrane to the cytoplasm is shown along with the diameter of the cytokinetic pore in microns. A schematic representation of the localization in the dividing NSC is shown below. (B) Quantification of Brat membrane release as a function of cytokinetic pore diameter for three different NSCs. (C) Microtubule dynamics during the late stages of NSC asymmetric division focusing on the cytokinetic pore and nascent NP sibling. Frames from Video 3 are shown with the microtubule marker Jupiter-GFP and the plasma membrane marker PLCδ-PH-mCherry (expressed with Worniu-GAL4 driven UAS). Time in minutes relative to central spindle compaction is shown along with the diameter of the cytokinetic pore in microns. A schematic representation of the localization in the dividing NSC is shown below. (D) Fascetto (Feo; PRC1 homolog) dynamics during the late stages of NSC asymmetric division focusing on the cytokinetic pore and nascent NP sibling. Frames from Video 3 are shown with Feo-GFP and the plasma membrane marker PLCδ-PH-mCherry (expressed with Worniu-GAL4 driven UAS). Time in minutes relative to Brat translocation from the membrane to the cytoplasm is shown along with the diameter of the cytokinetic pore in microns. A schematic representation of the localization in the dividing NSC is shown below. (E) Model for pre-division initiation of asymmetric fate specification. When mitosis is completed the nascent NSC and NP siblings remain connected by the intercellular bridge, which contains the recently-formed midbody. The midbody consists of central spindle microtubules (which extend between the divided nuclei) and other proteins such as Feo. Shortly after mitosis, fate determinants like Brat and Pros are released from the membrane but don’t exchange with the other nascent sibling’s cytoplasm. Pros is immediately imported into the nucleus of the nascent NP sibling following release.

### Pre-division fate determinant isolation requires the midbody

The ability of Brat and Pros to influence the fate of the NP sibling depends on their release from the plasma membrane. Our results indicate that the determinant activation that begins with membrane release occurs before cell division has completed. Furthermore, cytoplasmic exchange of the determinants between nascent siblings, which could corrupt the fate of the NSC sibling, is inhibited in late cytokinesis when determinants are released from the membrane. Release begins shortly after the cytokinetic pore reaches 1.5 µm in diameter, when the cytokinetic midbody forms. The midbody is a microtubule-rich structure that is constructed inside the pore formed by the intercellular bridge and facilitates abscission^31^. Furthermore, the midbody can inhibit exchange of proteins between nascent sibling cell cytoplasm before abscission^32^. We imaged microtubules and a marker of midbody microtubules, Fascetto (Feo; PRC1 in worms and mammals), which binds central spindle microtubules and bundles them during midbody formation, to examine midbody formation in asymmetric NSC divisions and verify that the midbody forms before fate determinant activation begins (Fig. 3C,D; Video 3). These data revealed that central spindle microtubules become compacted within the pore connecting the nascent siblings. The microtubules transition from having clear gaps during early furrowing, to a point at which there are no apparent gaps once the pore diameter reached approximately 1.5 µm (Fig. 3C,D; Video 3). Thus, the midbody forms a continuous structure in the pore nearly simultaneously with fate determinant membrane release. These observations are consistent with a potential role for the midbody in fate determinant activation by preventing determinant exchange between nascent siblings (Fig. 3E).

To determine if asymmetric fate specification requires cytoplasmic isolation by the midbody, we ablated the midbody shortly after formation and examined whether Brat was retained in the nascent NP cytoplasm. We used a cytoskeleton depolymerization strategy to ablate the midbody. In cultured cells and worm embryos, actin filament or microtubule depolymerization alone is insufficient to disrupt the midbody but their combined action ablate the structure^32,33^. Consistent with these reports, we found that addition of the actin depolymerizing drug Latrunculin A (LatA) to NSCs shortly before midbody formation caused furrow retraction, whereas depolymerization after formation inhibited further pore constriction while maintaining the connecting bridge (Fig. 4A,B; Video 4). We used Colcemid to depolymerize the midbody microtubules in LatA frozen midbodies and examined the dynamics of cytoplasmic Brat. Unlike in untreated or LatA alone-treated dividing NSCs, where Brat remained restricted to the cytoplasm of the nascent NP, we observed rapid Brat dissipation into the much larger NSC sibling in LatA + Colcemid treated cells (Fig. 4C-E; Video 4). We conclude that the midbody is required to maintain asymmetric distribution of fate determinants during late cytokinesis by acting as a barrier to cytoplasmic exchange between the sibling cells.

**Figure 4.**
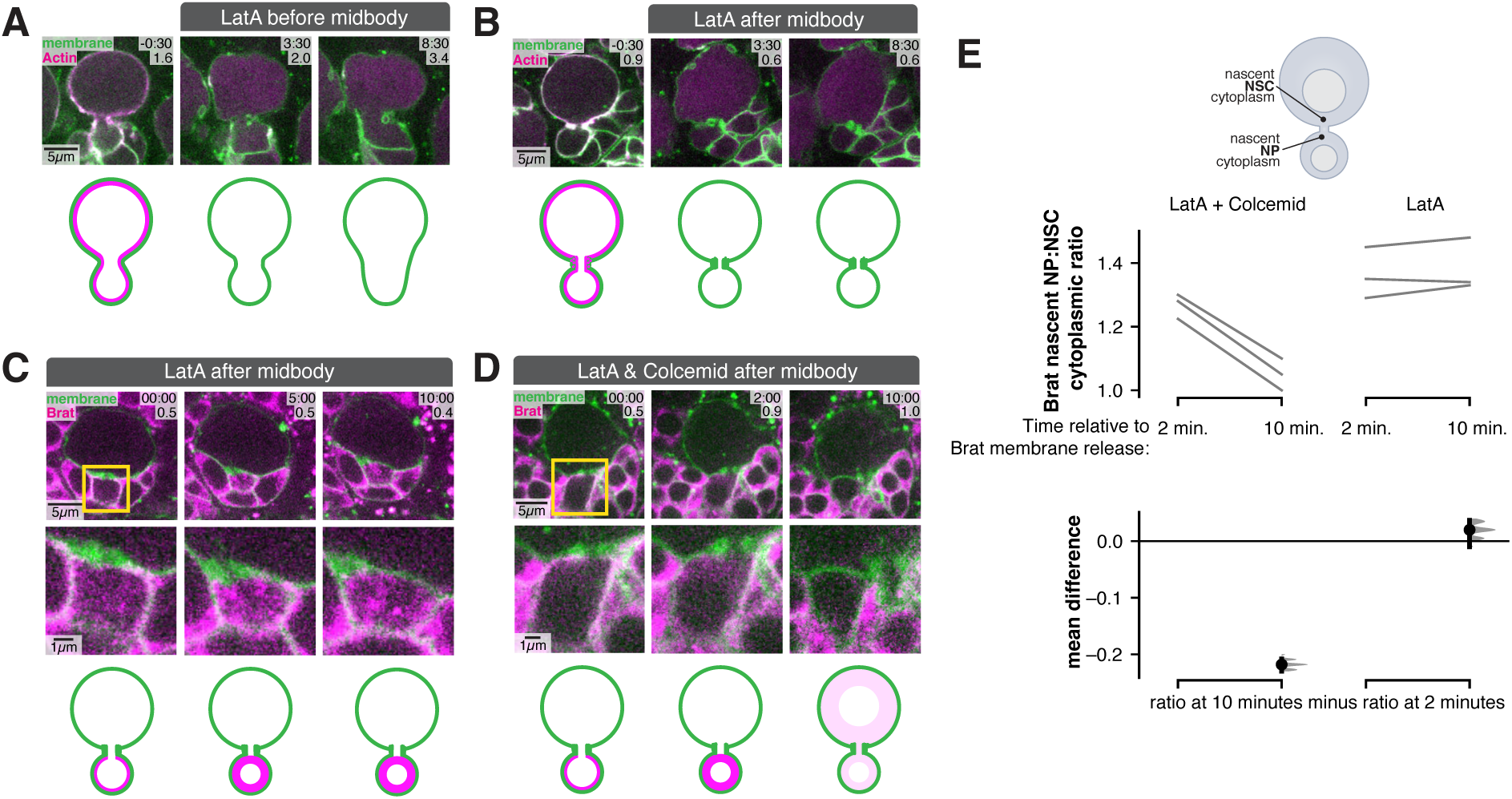
The cytokinetic midbody is required for cytoplasmic isolation during fate determinant activation. (A) Membrane and actin dynamics in NSC with the actin cytoskeleton depolymerized by LatA before midbody formation. Frames from Video 4 are shown with GMA-GFP (”Actin”) and PLCδ-PH-mCherry (both expressed with Worniu-GAL4 driven UAS). Time in minutes relative to Latrunculin A (LatA) addition is shown along with the diameter of the cytokinetic pore in microns. A schematized representation of the localization within the dividing NSC is shown below. (B) Membrane and actin dynamics in NSC with the actin cytoskeleton depolymerized by LatA after midbody formation. Frames from Video 4 are shown as in (A). (C) Brat dynamics in NSC with the actin cytoskeleton depolymerized by LatA before midbody formation. Frames from Video 4 are shown with Brat-GFP expressed from its endogenous locus and the plasma membrane marker PLCδ-PH-mCherry (expressed with Worniu-GAL4 driven UAS) through an optical section containing the cytokinetic pore. Time in minutes relative to Latrunculin A addition is shown along with the diameter of the cytokinetic pore in microns. Middle row shows inset of the cytokinetic pore and nascent NP sibling. A schematized representation of the localization within the dividing NSC is shown below. (D) Brat dynamics in NSC with midbody disrupted by LatA + Colcemid treatment. Frames from Video 4 are shown with Brat-GFP with time in minutes relative to LatA and Colcemid addition shown along with the diameter of the cytokinetic pore in microns. Middle row shows inset of the cytokinetic pore and nascent NP sibling. A schematized representation of the localization within the dividing NSC is shown below. (E) Quantification of the effect of actin and microtubule depolymerization on Brat asymmetry. The ratio of nascent NSC to NP cytoplasmic Brat signal is shown at 2 and 10 minutes after Brat membrane release. Each line represents paired measurements for an individual cell. Bar in the mean difference comparison represents bootstrap 95% confidence interval.

### Initiation of asymmetric fate specification requires midbody formation

Our results support a model in which the midbody plays a critical role in ACD by isolating nascent sibling cytoplasms from one another when fate determinants are released from the membrane, thus ensuring the fate of the incorrect sibling is not corrupted. The key role of the midbody in initiating asymmetric fate specification led us to ask if midbody formation could be a prerequisite for determinant membrane release. We tested this hypothesis by following Pros dynamics in NSCs in which midbody formation was inhibited. The microtubule stabilizing factor Fascetto (Feo; aka PRC1) is a key component of the midbody and is recruited to central spindle microtubules through interactions with the kinesin Klp3a (aka KIF4A)^34,35^. The interaction of Feo with Klp3a is promoted by Aurora B phosphorylation^36^. We verified that NSC midbody formation could be inhibited by addition of the Aurora B inhibitor Binucleine 2^37^. Dividing NSCs treated with inhibitor immediately before midbody formation lost Feo signal on the central spindle and failed to form a stable midbody, leading to furrow retraction (Fig. 5A; Video 5). In contrast, cells with recently formed midbodies maintained their furrows following Aurora B inhibition (Fig. 5B,C; Video 5). Nuclei from midbody inhibited NSCs imported a nuclear import marker at the completion of mitosis, indicating that exit from mitosis and subsequent import was not affected in these cells (Fig. 5D; Video 5). We examined Pros dynamics in Aurora B-inhibited NSCs, both before and after midbody formation. Pros release was significantly delayed in cells where midbody formation was inhibited by Aurora B (Fig. 5F,G). However, Pros was released from the membrane normally in Aurora B-inhibited cells that had already formed a midbody (Fig. 5E,G; Video 5). Consistent with the inhibition of membrane release, Pros also failed to enter the nucleus in midbody-inhibited NSCs (Fig. 5H). We also observed a requirement for the midbody for Brat membrane release (Fig. 5I-K). These results suggest that midbody formation is required for initiating asymmetric fate specification through release of fate determinants from the membrane.

**Figure 5.**
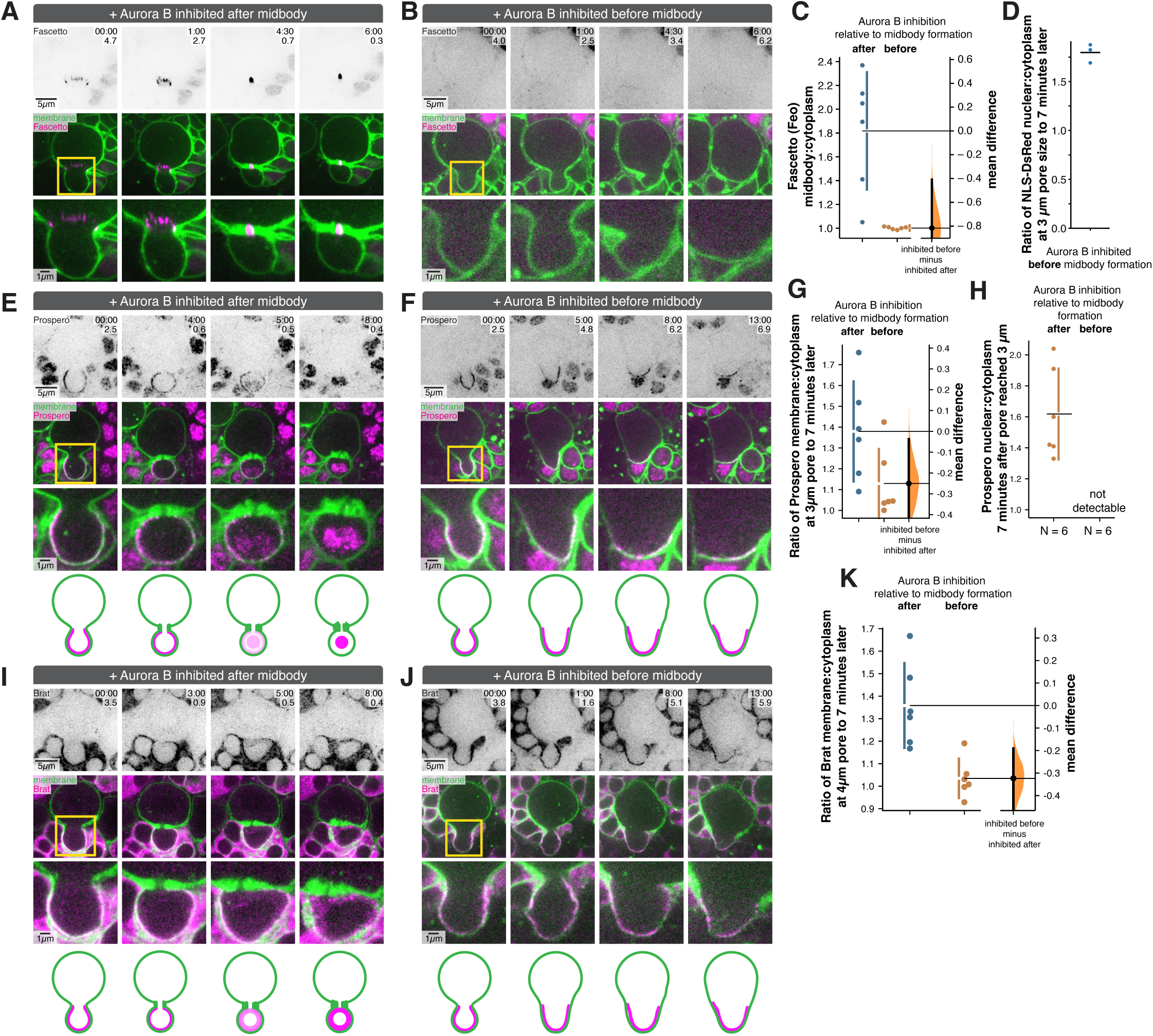
Fate determinant membrane release requires midbody formation. (A) Fascetto (Feo; aka PRC1) dynamics in NSC with Aurora B inhibited after midbody formation. Frames from Video 5 are shown with Feo-GFP expressed from the ubiquitin promoter and the plasma membrane marker PLCδ-PH-mCherry (expressed with Worniu-GAL4 driven UAS) through an optical section containing the cytokinetic pore in an NSC with Aurora B inhibited with the inhibitor Binucleine 2 immediately after the cytokinetic pore reached 1.5 µm in diameter. Time in minutes is shown along with the diameter of the cytokinetic pore in microns. Bottom row shows inset of the cytokinetic pore and nascent NP sibling. (B) Feo dynamics in NSC with Aurora B inhibited after midbody formation. Frames from Video 5 are shown as in (A) with Aurora B inhibited with the inhibitor Binucleine 2 immediately before the cytokinetic pore reaced 1.5 µm in diameter. (C) Quantification of Feo in NSCs with Aurora B inhibited before or after midbody formation. Gardner-Altman estimation plot of the ratio of Feo signal in the midbody to that of the cytoplasm is shown for six different NSC divisions with Aurora B inhibited before or after the cytokinetic pore reached 1.5 µm. The error bars represent one standard deviation (gap is mean); the bar in the mean difference comparison represents bootstrap 95% confidence interval. (D) Quantification of nuclear import in NSCs with Aurora B inhibited before midbody formation. The ratio of nuclear:cytoplasmic Nuclear Localization Signal-DsRed (NLS-DsRed) at a pore size of 3 µM to seven minutes afterwards is shown. (E) Prospero (Pros) dynamics in NSC with Aurora B inhibited after midbody formation. Frames from Video 5 are shown with Pros-GFP expressed from its endogenous locus and the plasma membrane marker PLCδ-PH-mCherry (expressed with Worniu-GAL4 driven UAS) through an optical section containing the cytokinetic pore in an NSC with Aurora B inhibited with the inhibitor binucleine immediately after the cytokinetic pore reached 1.5 µm in diameter. Time in minutes is shown along with the diameter of the cytokinetic pore in microns. An inset of the cytokinetic pore and nascent NP sibling is shown. The bottom row is a schematized representation of the localization within the dividing NSC. (F) Pros dynamics in NSC with Aurora B inhibited before midbody formation. Frames from Video 5 are shown as in (E) with Aurora B inhibited with the inhibitor Binucleine 2 immediately before the cytokinetic pore reaced 1.5 µm in diameter. (G) Quantification of Pros membrane dynamics in dividing NSCs when Aurora B was inhibited before or after midbody formation. Gardner-Altman estimation plot of the membrane to cytoplasmic Pros signal in the nascent NP when the cytokinetic pore reached 3 µm divided by membrane to cytoplasmic Pros seven minutes afterwards. The error bars represent one standard deviation (gap is mean); the bar in the mean difference comparison represents bootstrap 95% confidence interval. (H) Quantification of Pros nuclear dynamics in dividing NSCs when Aurora B was inhibited before or after midbody formation. The ratio of nuclear to cytoplasmic Pros signal in the nascent NP is shown seven minutes following when the cytokinetic pore reached 3 µm for NSCs with Aurora B inhibited with the inhibitor Binucleine 2 immediately before or after the cytokinetic pore reached 1.5 µm in diameter. The error bar represents one standard deviation (gap is mean). (I) Brain Tumor (Brat) dynamics in NSC with Aurora B inhibited after midbody formation. Frames from Video 5 are shown with Brat-GFP expressed from its endogenous locus and the plasma membrane marker PLCδ-PH-mCherry (expressed with Worniu-GAL4 driven UAS) through an optical section containing the cytokinetic pore in an NSC with Aurora B inhibited with the inhibitor Binucleine 2 immediately after the cytokinetic pore reached 1.5 µm in diameter. Time in minutes is shown along with the diameter of the cytokinetic pore in microns. An inset of the cytokinetic pore and nascent NP sibling is shown. The bottom row is a schematized representation of the localization within the dividing NSC. (J) Brat dynamics in NSC with Aurora B inhibited before midbody formation. Frames from Video 5 are shown as in (I) with Aurora B inhibited with the inhibitor Binucleine 2 immediately before the cytokinetic pore reaced 1.5 µm in diameter. (K) Quantification of Brat membrane dynamics in dividing NSCs when Aurora B was inhibited before or after midbody formation. Gardner-Altman estimation plot of the membrane to cytoplasmic Brat signal in the nascent NP when the cytokinetic pore reached 4 µm divided by membrane to cytoplasmic Brat seven minutes afterwards. The error bars represent one standard deviation (gap is mean); the bar in the mean difference comparison represents bootstrap 95% confidence interval.

### The cell cycle phosphatase String regulates fate determinant release after midbody formation

Our results indicate that the release of fate determinants from the membrane that initiates asymmetric fate specification occurs before cell division completes. Formation of the midbody appears to be a prerequisite for determinant membrane release such that the signals that control release appear to be activated only after the midbody isolates nascent sibling cytoplasms from one another. We sought to identify signaling factors that regulate release of determinants from the membrane post-midbody formation and screened cell cycle inhibitors for their effect on Pros membrane release attempting to identify regulators that promote release without disrupting midbody formation. Addition of an inhibitor of String (Cdc25 in mammals), a cell cycle phosphatase, after the onset of furrowing significantly delayed Pros membrane release without any detectable effect on midbody formation (Fig. 6A,B; Video 6). Although Cdc25 has been reported to be degraded near the end of mitosis, we detected signal with an anti-String antibody in NSCs during late cytokinesis (Fig. 6C).

**Figure 6.**
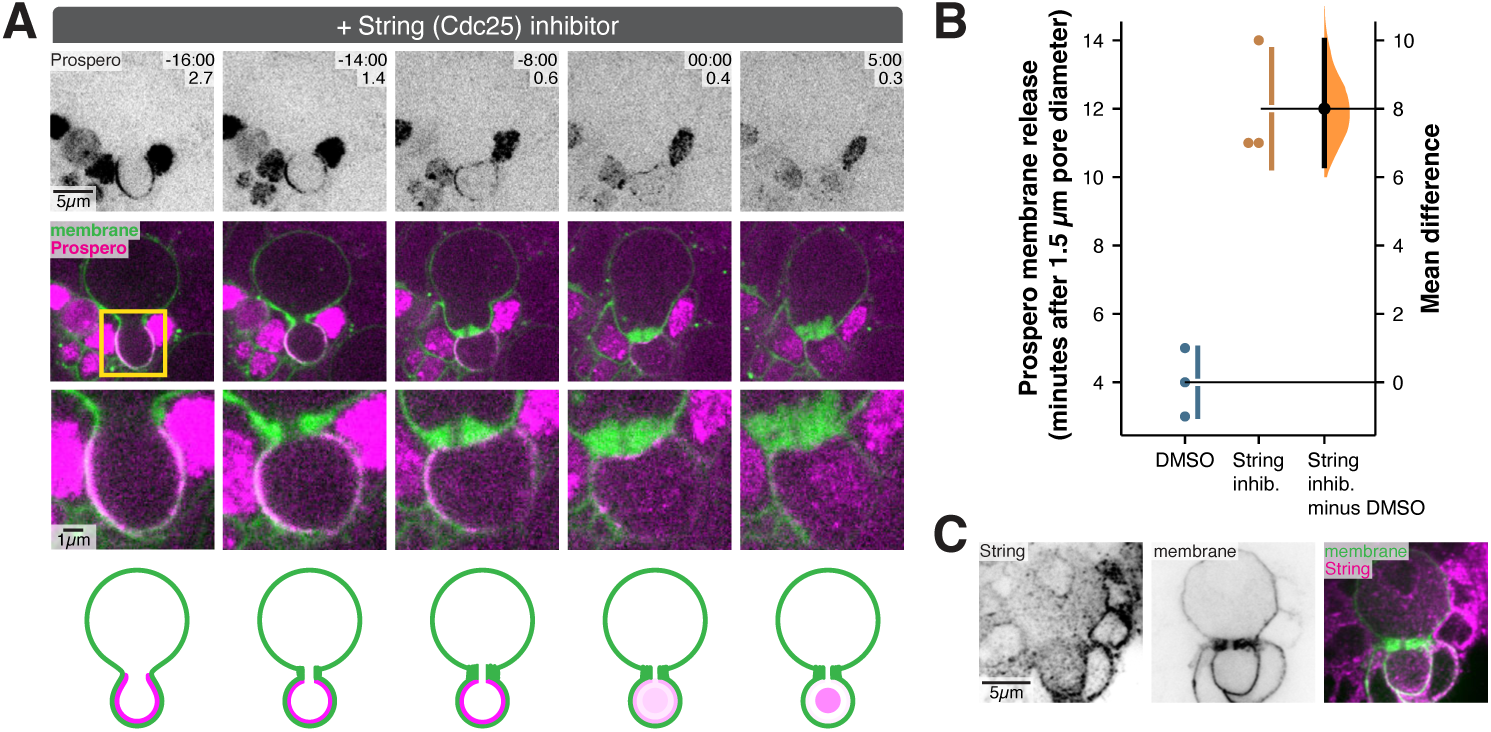
The cell cycle phosphatase String (Cdc25) is required for fate determinant membrane release following midbody formation. (A) Inhibition of String (Cdc25) leads to delayed Pros release. Frames from Video 6 are shown with Pros-GFP expressed from its endogenous locus and the plasma membrane marker PLCδ-PH-mCherry (expressed with Worniu-GAL4 driven UAS) through an optical section containing the cytokinetic pore in NSCs treated with the String inhibitor. Time in minutes relative to the Pros membrane release is shown along with the diameter of the cytokinetic pore in microns. Merge is shown in the second row along with an inset in the third row that focuses on the cytokinetic pore and NP sibling. Bottom row shows a schematized representation of the localization. (B) Quantification of the effect of String inhibition on Pros membrane release. Gardner-Altman estimation plot of the number of minutes after the cytokinetic pore reached 1.5 µm that Pros was released from the membrane. The error bars represent one standard deviation (gap is mean); the bar in the mean difference comparison represents bootstrap 95% confidence interval. (C) String localization during late NSC cytokinesis. An NSC expressing PLCδ-PH-GFP (expressed with Worniu-GAL4 driven UAS) fixed and stained with anti-String (”String”) and anti-GFP (”membrane”) antibodies is shown.

## Discussion

We examined fate determinant dynamics at high temporal resolution and discovered a new function for the cytokinetic midbody in ACD. This function is essential for timely fate determinant activation, highlighting the importance of determinant regulation during ACD. Fate determinants such as Brat and Pros are polarized on the plasma membrane during division as a mechanism for their segregation, but it was not known when they are released from the membrane relative to both nuclear and cell division. We hypothesized that they would not be released until sometime after division to ensure that they do not enter the cytoplasm of the incorrect sibling cell, potentially corrupting its fate. Surprisingly, however, we found that asymmetric fate specification is initiated by fate determinant membrane release long before cell division completes. Determinants are rapidly released after nuclear division, while the nascent sibling cells remain attached by the midbody-containing intercellular bridge. Brat and Pros accumulate in the nascent NP cytoplasm and nucleus, respectively, without exchanging into the nascent NSC, indicating that determinants do not exchange between the siblings. Disrupting the midbody causes Brat to leak out of the nascent NP indicating that the midbody is required to prevent determinant diffusion between the nascent siblings. These results provide a framework for understanding determinant activation and support a key role for the midbody in ACD.

The midbody was discovered in the late 19^th^ century by early cell division researchers^38^ and its highly organized structure suggests an important role in cellular function. Only recently, this was borne out by the discovery that the midbody recruits and positions the abscission machinery once the pore has constricted to a diameter of 200 nm^24^. Our discovery suggests that the midbody is important at a much earlier timepoint, shortly after it is formed from the compacted central spindle and associated proteins. Midbody-connected siblings can persist well into the following interphase^39^, forming a temporal window between mitotic and cytokinetic exit. Taking advantage of this period to specify cell state could be important in rapidly proliferating tissues like the developing brain. Thus, the midbody allows asymmetric fate specification to begin immediately following nuclear division, when large-scale transcriptional activity has resumed.

The barrier function of the NSC midbody is reminiscent of the bud neck in budding yeast by acting as a diffusion barrier^27,28^. However, not all midbodies appear to prevent exchange across the intercellular bridge. An early study found that molecules as large as antibodies pass across the bridge formed by dividing HeLa cells^40^. More recently cultured mouse embryonic stem cells have been demonstrated to remain connected by an freely passable intercellular bridge following mitosis, only exiting naïve pluripotency after division^12^. Other midbodies, besides the NSC’s, restrict passage, however. The midbody formed by the first division of the worm zygote prevents exchange^32^, as does the meiotic midbody formed by mammalian oocytes^41^. This diversity suggests that the midbody structure might have the capability to be tuned to inhibit or allow diffusion between nascent siblings.

We propose that fate determinant membrane release is part of a highly choreographed sequence of events at the end of mitosis and during late cytokinesis that are centered around the midbody. Midbody formation, characterized by central spindle compaction and straightening of the furrow membrane into the highly cylindrical intercellular bridge, takes place alongside mitotic exit (Figs. 2,3; Videos 2 & 3). The pore constricts from approximately 1.5 µm to 200 nm, the diameter at which abscission can occur, and fate determinants release from the membrane shortly after constriction begins. Our results suggest that the mitotic phosphatase String is required for Pros membrane release after midbody formation. Further work will be required to understand how String activity might be connected to midbody dynamics to ensure that Pros is not released before the cytoplasmic exchange barrier has been established.

## Resource Availability

### Lead Contact

Contact the Lead Contact, Kenneth Prehoda, for further information or to request resources and reagents.

## Materials Availability

No new reagents were generated in this study.

## Data and Code Availability

Raw data available from the corresponding author on request.

## Experimental Model and Subject Details

### Fly Strains

Tissue specific expression of UAS controlled transgenes in NSCs was achieved using a Worniu-GAL4 driver line. Membrane dynamics were imaged using the membrane markers UAS-PLCδ-PH-GFP and UAS-PLCδ-PH-mCherry, which express the pleckstrin homology domain of human PLCδ tagged with GFP or mCherry, and binds to the plasma membrane lipid phosphoinositide PI(4,5)P_2_. F-Actin was visualized using UAS-GMA-GFP, which expresses a GFP tagged actin binding domain of Moesin. The onset of nuclear import at the end of mitosis was monitored using UAS-NLS-DsRed, which is comprised of a DsRed protein containing a nuclear localization signal (NLS) and expressed under the control of UAS.

Microtubules were imaged using GFP tagged Jupiter^42^. Fascetto (Feo) was imaged using a GFP tagged Fascetto protein under control of ubiquitin regulatory sequences. GFP tagged Prospero (Pros), Brain Tumor (Brat), and Deadpan (Dpn) proteins were generated by the modERN Project^43^. The nuclear envelope was imaged using GFP tagged Klaroid^44^. Chromosomes were imaged using RFP tagged Histone 2A (His2a).

## Method Details Live Imaging

To obtain brain explants, third instar *Drosophila* larvae were dissected in Schneider’s Insect Media (SIM) and the central nervous system was isolated. Next, larval brain explants were mounted on sterile poly-D-lysine coated 35mm glass bottom dish (ibidi Cat#81156) containing modified minimal hemolymph-like solution (HL3.1). Then, brain explants were imaged using a Nikon Eclipse Ti-2 Yokogawa CSU-W1 SoRa spinning disk microscope equipped dual Photometrics Prime BSI sCMOS cameras using a 60x H_2_O objective. 488 nm light was used to illuminate GFP tagged proteins and 561 nm light was used to illuminate DsRed and mCherry tagged proteins. Super resolution imaging was achieved by using SoRa (super resolution through optical photon reassignment) optics^45^. NSCs were identified by their large size, location in the central nervous system, and the use of NSC specific tissue driver lines. Time lapse imaging of midbody dynamics was achieved by refocusing the imaging plane on the medial plane of the cleavage furrow, and subsequently the midbody, along the apical-basal axis just before capturing each frame. Pharmacological inhibition of Aurora B was performed using 15 μM Binucleine 2 solubilized in DMSO. Pharmacological depolymerization of microtubules was performed using 1 mM Colcemid solubilized in DMSO. Pharmacological depolymerization of F-actin was performed using 50 μM Latrunculin A (LatA) solubilized in DMSO. Pharmacological inhibition of String was performed using 750 μM of the Cdc25 inhibitor NSC 663284 solubilized in DMSO.

## Immunofluorescence Staining

To determine the localization of native String in NSCs, the central nervous systems of third instar *Drosophila* larvae expressing Worniu-GAL4>UAS-PLCδ-PH-GFP were fixed in 4% paraformaldehyde. Mouse anti-GFP antibodies were used to stain NSC membranes (marked by Worniu-GAL4 driven UAS-PLCδ-PH-GFP), and guinea pig anti-String antibodies were used to stain String. Primary antibodies were used at a concentration of 1:100 (anti-GFP) and 1:75 (anti-String). Alexa 488 labeled anti-mouse (Invitrogen) and Cy3 labeled anti-guinea pig (Jackson Labs) secondary antibodies were used at a concentration of 1:500. Super resolution images were captured using a Nikon Eclipse Ti-2 Yokogawa CSU-W1 SoRa spinning disk microscope equipped dual Photometrics Prime BSI sCMOS cameras using a 60x H2O objective.

## Image Processing and Analysis

Imaging data was processed using ImageJ (FIJI package). For some movies, the bleach correction tool was used to correct for photobleaching. To reduce noise in Deadpan images, Gaussian blur was applied. For quantifying the dynamics of the cytokinetic pore size, medial sections (along the apical-basal axis) were used to measure the width of the cytokinetic pore. If ever the pore moved out of the focal plane, the pore size of the previous frame was used.

Quantification for Fig. 1D: The cytokinetic pore diameter was measured at the onset of Prospero membrane release for n=10 dividing NSCs.

Quantification for Fig. 1E: To quantify cytokinetic pore closure dynamics, the pore size was measure as a function of time and plotted for n=5 dividing NSCs.

Quantification for Fig. 1G: The cytokinetic pore diameter was measured at the onset of Deadpan nuclear import for n=3 dividing NSCs.

Quantification for Fig. 2B: The cytokinetic pore diameter was measured at the onset of chromosome decompaction, nuclear separation, and the onset of nuclear import. His2a-RFP signal was used to determine when chromosome began to decompact for n=3 dividing NSCs. Klaroid-GFP was used to determine when the nascent sibling nuclear compartments became separated for n=3 dividing NSCs. NLS-DsRed was used to determine the onset of nuclear import for n=3 dividing NSCs. Nuclear import was determined by the frame in which nuclear intensity of NLS-DsRed began to increase.

Quantification for Fig. 3B: To quantify the timing of Brat membrane release during NSC division, Image J was used to measure average cytoplasmic signal intensity in the nascent NP and nascent NSC sibling cytoplasms. The ratio of nascent NP:NSC cytoplasmic Brat signal was plotted as a function of cytokinetic pore constriction.

Quantification for Fig. 4E: To quantify the effect of actin and microtubule depolymerization on Brat asymmetry, Brat signal in the nascent NSC and NP cytoplasms was measured to calculate the nascent NP:NSC cytoplasmic Brat ratio. This ratio was calculated at 2 and 10 minutes after Brat membrane release was detected. Latrunculin A treated dividing NSCs (n=3) were compared to Latrunculin A + Colcemid treated dividing NSCs (n=3).

Quantification for Fig. 5C: To quantify the effect of Aurora B inhibition on Fascetto dynamics, Fascetto signal intensity within the midbody and within the nascent NSC cytoplasm was measured to calculate a midbody:cytoplasmic ratio. Measurements were taken when the cytokinetic pore constricted to under 3 µm. Using Image J, average signal intensity within a 3 µm x 3 µm box drawn over the midbody was used to measure Fascetto signal intensity in the midbody. The same size box was used to measure average Fascetto signal intensity within the nascent NSC cytoplasm adjacent to the midbody. Dividing NSCs where Aurora B inhibition occurred after midbody formation (n=6) were compared to dividing NSCs where Aurora B inhibition occurred before midbody formation (n=6).

Quantification for Fig. 5D: To determine the effect of Aurora B inhibition on nuclear import dynamics, nuclear:cytoplasmic ratio of the nuclear marker NLS-DsRed was calculated for dividing NSCs where Aurora B inhibition occurred before midbody formation. Average signal intensity of NLS-DsRed within the nuclear compartment and within the cytoplasm was measured to calculate a nuclear:cytoplasm ratio for n=3 dividing NSCs.

Quantification for Fig. 5G: To quantify the effect of Aurora B inhibition on Prospero dynamics, Prospero signal intensity on the nascent NP membrane and within the nascent NP cytoplasm was measured to calculate a membrane:cytoplasm ratio. Using Image J, Prospero membrane intensity was measured by tracing the plasma membrane (marked by UAS-PLCδ-PH-mCherry) and then copying the ROI (region of interest) to the Prospero channel. A line of similar length was drawn within the nascent NP cytoplasm. Measurements were taken when the cytokinetic pore constricted to under 3 µm and then again 7 minutes later. Dividing NSCs where Aurora B inhibition occurred after midbody formation (n=6) were compared to dividing NSCs where Aurora B inhibition occurred before midbody formation (n=6).

Quantification for Fig. 5H: Nuclear Prospero signal intensity was measured by outlining the nascent NP nucleus and measuring average signal intensity within. Nuclear:cytoplasm ratios were calculared 7 minutes after the pore has constricted down to 3 µm. Dividing NSCs where Aurora B inhibition occurred after midbody formation (n=6) were compared to dividing NSCs where Aurora B inhibition occurred before midbody formation (n=6).

Quantification for Fig. 5K: To quantify the effect of Aurora B inhibition on Brat dynamics, Brat signal intensity on the nascent NP membrane and within the nascent NP cytoplasm was measured to calculate a membrane:cytoplasm ratio. Using Image J, Brat membrane intensity was measured by tracing the plasma membrane (marked by UAS-PLCδ-PH-mCherry) and then copying the ROI (region of interest) to the Brat channel. A line of similar length was drawn within the nascent NP cytoplasm. Measurements were taken when the cytokinetic pore constricted to under 4 µm and then again 7 minutes later. Dividing NSCs where Aurora B inhibition occurred after midbody formation (n=6) were compared to dividing NSCs where Aurora B inhibition occurred before midbody formation (n=6).

Quantification for Fig. 6B: The effect of String inhibition on Prospero dynamics was measured by, waiting until the cytokinetic pore constricted to 1.5 µm and then determining the time until onset Prospero nuclear import into the nascent NP nucleus. Dividing NSCs treated with the String inhibitor NSC 663284 (n=3) were compared to DMSO alone (n=3).

## Statistical Analysis

Gardner-Altman estimation plots and 95% confidence intervals of datasets were prepared using the DABEST package^46^. Statistical details can be found in the relevant methods section and figure legend

Key Resources Table.

Key Resources Table
| REAGENT or RESOURCE | SOURCE | IDENTIFIER |
| --- | --- | --- |
| **Experimental Model: Fly Strains** |  |  |
| Brain Tumor (Brat)-GFP | Bloomington *Drosophila* Stock Center (BDSC) | BDSC Cat#83658<br>RRID: BDSC 83658 |
| Deadpan (Dpn)-GFP | Bloomington *Drosophila* Stock Center (BDSC) | BDSC Cat#65295<br>RRID: BDSC 65295 |
| Fascetto (Feo)-GFP | Bloomington *Drosophila* Stock Center (BDSC) | BDSC Cat#59273<br>RRID: BDSC 59273 |
| His2a-RFP<br>(chromosome marker) | Bloomington *Drosophila* Stock Center (BDSC) | BDSC Cat# 23650<br>RRID: BDSC 23650 |
| Jupiter-GFP<br>(microtubule marker) | Bloomington *Drosophila* Stock Center (BDSC) | BDSC Cat#60156<br>RRID: BDSC 60156 |
| Klaroid-GFP<br>(nuclear envelope marker) | Bloomington *Drosophila* Stock Center (BDSC) | BDSC Cat#51525<br>RRID: BDSC 51525 |
| Prospero (Pros)-GFP | Bloomington *Drosophila* Stock Center (BDSC) | BDSC Cat#66463<br>RRID: BDSC 66463 |
| UAS-GMA-GFP<br>(F-actin marker) | Bloomington *Drosophila* Stock Center (BDSC) | BDSC Cat#31776<br>RRID: BDSC 31776 |
| UAS-PLCδ-PH-GFP<br>(membrane marker) | Bloomington *Drosophila* Stock Center (BDSC) | BDSC Cat#39693<br>RRID: BDSC 39693 |
| UAS-PLCδ-PH-mCherry<br>(membrane marker) | Bloomington *Drosophila* Stock Center (BDSC) | BDSC Cat#51658<br>RRID: BDSC 51658 |
| UAS-NLS-DsRed<br>(nuclear marker) | Bloomington *Drosophila* Stock Center (BDSC) | BDSC Cat#8547<br>RRID: BDSC 8547 |
| Worniu-GAL4<br>(NSC driver line) | Chris Q. Doe Lab |  |
| **Pharmacological Inhibitors** |  |  |
| Binucleine 2 | Sigma-Aldrich | Cat#B1186 |
| Colcemid | Millipore | Cat#234109-M |
| Latrunculin A (LatA) | Enzo | Cat#BML-T119-0100 |
| NSC 663284 | Cayman Chemical | Cat#13303 |
| **Primary Antibodies** |  |  |
| anti-GFP | Developmental Studies | Cat# DSHB-GFP-12A6 |
|  | Hybridoma Bank | RRID: AB_2617417 |
| anti-String | Yukiko M. Yamashita Lab |  |

## Video Legends

**Video 1: Asymmetric fate specification is initiated before NSC division completes**

Part 1: Prospero dynamics during the late stages of NSC asymmetric division. Super resolution videos of NSCs expressing Prospero-GFP “Prospero” and the membrane marker UAS-PLCδ-PH-mCherry “membrane”. Time relative to start of nuclear import of Prospero is indicated. Three movies of dividing NSCs are shown.

Part 2: Deadpan dynamics during the late stages of NSC asymmetric division. Super resolution videos of NSCs expressing Deadpan-GFP “Deadpan” and the membrane marker UAS-PLCδ-PH-mCherry “membrane”. To improve image quality, Gaussian blur was applied to the Deadpan channel. Time relative to start of nuclear import of Deadpan is indicated. Three movies of dividing NSCs are shown.

**Video 2: A temporal window between mitosis and cytokinesis during NSC asymmetric cell division**

Part 1: Chromatin dynamics during late NSC division. Super resolution videos of NSCs expressing the chromosome marker His2a-RFP “His2a” and the membrane marker UAS-PLCδ-PH-GFP “membrane”. Time relative to start of chromosome decompaction is indicated. Three movies of dividing NSCs are shown.

Part 2: Nuclear dynamics during late NSC division. Super resolution videos of NSCs expressing the nuclear envelope marker Klaroid-GFP “Klaroid” and the membrane marker UAS-PLCδ-PH-mCherry “membrane”. Time relative to nuclear membrane separation. Three movies of dividing NSCs are shown.

Part 3: Nuclear import dynamics during late NSC division. Super resolution videos of NSCs expressing the nuclear marker UAS-NLS-DsRed “NLS-DsRed” and the membrane marker UAS-PLCδ-PH-GFP “membrane”. Time relative to start of nuclear import is indicated. Three movies of dividing NSCs are shown.

**Video 3: Fate determinants are isolated in nascent siblings before NSC division completes**

Part 1: Brat dynamics during late NSC division. Super resolution videos of NSCs expressing Brain Tumor (Brat)-GFP “Brat” and the membrane marker UAS-PLCδ-PH-mCherry “membrane”. Time relative to membrane release of Brat. Three movies of dividing NSCs are shown.

Part 2: Microtubule dynamics during late NSC division. Super resolution videos of NSCs expressing the microtubule marker Jupiter-GFP “microtubules” and the membrane marker UAS-PLCδ-PH-mCherry “membrane”. Bottom row is a zoomed-in view of the cytokinetic pore. Time relative to midbody formation. Three movies of dividing NSCs are shown.

Part 3: Fascetto dynamics during late NSC division. Super resolution videos of NSCs expressing Fascetto-GFP “Fascetto” and the membrane marker UAS-PLCδ-PH-mCherry “membrane”. Bottom row is a zoomed-in view of the cytokinetic pore. Time relative to midbody formation. Three movies of dividing NSCs are shown.

**Video 4: The cytokinetic midbody is required for cytoplasmic isolation during fate determinant activation**

Part 1: Membrane and actin dynamics in NSC with the actin cytoskeleton depolymerized by LatA before midbody formation. Super resolution videos of NSCs expressing the F-actin marker UAS-GMA-GFP “F-actin” and the membrane marker UAS-PLCδ-PH-mCherry “membrane” in the presence of the F-actin inhibitor, Latrunculin A (LatA). Bottom row is a zoomed-in view of the cytokinetic pore. Time relative to LatA addition.

Part 2: Membrane and actin dynamics in NSC with the actin cytoskeleton depolymerized by LatA after midbody formation. Super resolution videos of NSCs expressing the F-actin marker UAS-GMA-GFP “F-actin” and the membrane marker UAS-PLCδ-PH-mCherry “membrane” in the presence of the F-actin inhibitor, Latrunculin A (LatA). Bottom row is a zoomed-in view of the cytokinetic pore. Time relative to LatA addition.

Part 3: Brat dynamics in NSCs with the actin cytoskeleton depolymerized by LatA after midbody formation. Super resolution videos of NSCs expressing Brain Tumor (Brat)-GFP “Brat” and the membrane marker UAS-PLCδ-PH-mCherry “membrane” in the presence of the F-actin inhibitor, Latrunculin A (LatA). The drug was added just after midbody formation. Time relative to membrane release of Brat. Three movies of dividing NSCs are shown.

Part 4: Brat dynamics in NSC with midbody disrupted by LatA + Colcemid treatment. Super resolution videos of NSCs expressing Brain Tumor (Brat)-GFP “Brat” and the membrane marker UAS-PLCδ-PH-mCherry “membrane” in the presence of the F-actin inhibitor, Latrunculin A (LatA), and the microtubule inhibitor, Colcemid. The drug cocktail was added just after midbody formation. Time relative to membrane release of Brat. Three movies of dividing NSCs are shown.

**Video 5: Fate determinant membrane release requires midbody formation**

Part 1: Fascetto dynamics in NSCs with Aurora B inhibited before midbody formation. Super resolution videos of NSCs expressing Fascetto-GFP “Fascetto” and the membrane marker UAS-PLCδ-PH-mCherry “membrane” in the presence of the Aurora B inhibitor, Binucleine 2. Time relative to start of imaging. Six movies of dividing NSCs are shown.

Part 2: Fascetto dynamics in NSCs with Aurora B inhibited after midbody formation. Super resolution videos of NSCs expressing Fascetto-GFP “Fascetto” and the membrane marker UAS-PLCδ-PH-mCherry “membrane” in the presence of the Aurora B inhibitor, Binucleine 2. Time relative to start of imaging. Six movies of dividing NSCs are shown.

Part 3: Prospero dynamics in NSCs with Aurora B inhibited before midbody formation. Super resolution videos of NSCs expressing Prospero-GFP “Prospero” and the membrane marker UAS-PLCδ-PH-mCherry “membrane” in the presence of the Aurora B inhibitor, Binucleine 2. Time relative to start of imaging. Six movies of dividing NSCs are shown.

Part 4: Prospero dynamics in NSCs with Aurora B inhibited after midbody formation. Super resolution videos of NSCs expressing Prospero-GFP “Prospero” and the membrane marker UAS-PLCδ-PH-mCherry “membrane” in the presence of the Aurora B inhibitor, Binucleine 2Time relative to start of imaging. Six movies of dividing NSCs are shown.

Part 5: Brat dynamics in NSCs with Aurora B inhibited before midbody formation. Super resolution videos of NSCs expressing Brain Tumor (Brat)-GFP “Brat” and the membrane marker UAS-PLCδ-PH-mCherry “membrane” in the presence of the Aurora B inhibitor, Binucleine 2. Time relative to start of imaging. Six movies of dividing NSCs are shown.

Part 6: Brat dynamics in NSCs with Aurora B inhibited after midbody formation. Super resolution videos of NSCs expressing Brain Tumor (Brat)-GFP “Brat” and the membrane marker UAS-PLCδ-PH-mCherry “membrane” in the presence of the Aurora B inhibitor, Binucleine 2. Time relative to start of imaging. Six movies of dividing NSCs are shown.

Part 7: Nuclear import dynamics in NSCs with Aurora B inhibited before midbody formation. Super resolution videos of NSCs expressing the nuclear marker UAS-NLS-DsRed “NLS-DsRed” and the membrane marker UAS-PLCδ-PH-GFP “membrane in the presence of the Aurora B inhibitor, Binucleine 2. Time relative to start of imaging. Three movies of dividing NSCs are shown.

**Video 6: The cell cycle phosphatase String (Cdc25) is required for fate determinant membrane release following midbody formation**

Part 1: Prospero dynamics with String (Cdc25) inhibited. Super resolution videos of NSCs expressing Prospero-GFP “Prospero” and the membrane marker UAS-PLCδ-PH-mCherry “membrane” in the presence of the String inhibitor, NSC 663284. The drug was added just prior to midbody formation. Time relative to start of nuclear import of Prospero is indicated. Three movies of dividing NSCs are shown.

Part 2: Prospero dynamics in NSCs treated with DMSO (vehicle control). Prospero-GFP “Prospero” and the membrane marker UAS-PLCδ-PH-mCherry “membrane” in the presence of 2% DMSO. The DMSO was added just prior to midbody formation. Time relative to start of nuclear import of Prospero is indicated. Three movies of dividing NSCs are shown.

## Acknowledgments

We thank Brad Nolen and Daniel Grimes for their useful comments on this manuscript. We thank Adam Fries for maintaining the microscope used in this study. We thank Yukiko Yamashita for the anti-String antibody. This work was supported by NIH grants R35GM127092 and K99GM147601.

## Author Contributions

B.L. and K.E.P. designed the experiments. B.L. performed the experiments. R.R.P. contributed to the String inhibition and String staining experiments. B.L. and K.E.P analyzed the data, prepared the figures, and wrote the manuscript.

